# Influence of High Concentrations of Copper Sulfate on *In Vitro* Adventitious Organogenesis of *Cucumis sativus* L.

**DOI:** 10.1101/2021.02.24.432794

**Authors:** Jorge Fonseca Miguel

## Abstract

The effects of different concentrations of copper sulfate (0.2 to 5 mg L^−1^) on *in vitro* callus and shoot formation of cucumber was investigated. Four-day-old cotyledon explants from the inbred line ‘Wisconsin 2843’ and the commercial cultivars ‘Marketer’ and ‘Negrito’ were used. The results on callus-derived shoots showed that the optimal concentration of CuSO_4_ added to Murashige & Skoog (MS)-derived shoot induction medium containing 0.5 mg L^−1^ IAA and 2.5 mg L^−1^ BAP was 8-200 fold greater than in standard MS medium, and was genotype dependent. The highest genotypes response on shoot frequency and shoot number was achieved in this order by ‘Marketer’, ‘Negrito’ and ‘Wisconsin 2843’ with 1, 0.2 y 5 mg L^−1^ CuSO_4_, respectively. The genotype with the lowest control performance demanded the highest concentration of CuSO_4_ for its optimal morphogenic response - 6- and 10-fold more in shoot frequency and shoot number, respectively. The other cultivars registered a 2-fold increase in both variables. All explants formed callus and the response on callus extension varied among cultivars. The regression analysis showed a statistically significant relationship between shoot number and concentrations of CuSO_4_ and absence of association with callus extension. The present results indicate that application of specific concentrations of CuSO_4_ higher than in standard MS medium, increases adventitious cucumber shoot organogenesis.

## 1. Introduction

Cucumber (*Cucumis sativus* L.) belongs to the Cucurbitaceae family. This species is an economically important vegetable crop and a model plant for sex determination and vascular biology studies (Huang et al., 2009). According to the Food and Agriculture Organization of the United Nations (FAO), in 2019, the world production of cucumbers, including gherkins, ranked third among vegetable crops with 87,805,086 tonnes, and a harvested area of 2,231,402 hectares (FAOSTAT, http://www.fao.org/faostat/en/#data/QC). Crop improvement of cucumber for resistance/tolerance against major biotic and abiotic stresses is difficult by conventional breeding due to its narrow genetic base, a genetic variability of only 3-8% (Plader et al., 2007), and several crossing barriers with related species (den Nijs and Custers, 1990). Applying genetic engineering and plant transformation technologies make it possible to overcome these limitations and to develop cultivars with value-added traits.

Despite the number of publications on genetic transformation of cucumber (for reviews, see Yin et al., 2005; He et al., 2008; Wang et al, 2015; Miguel, 2017), with *Agrobacterium*-mediated gene transfer as the most commonly used, and reliable method, its efficiency is still very low (Wang et al., 2015; Miguel, 2017). One of the main problems in obtaining transgenic cucumber plants is related to regeneration, i.e. low morphogenic response in explants of certain genotypes, and decrease in the regeneration rate as a result of the usual transformation steps (Miguel, 2017). A reliable plant genetic transformation system demands an efficient and stable regeneration procedure (Yin et al., 2005; He et al., 2008; Wang et al., 2015). In almost all practical transformation systems, tissue culture is employed to achieve a workable efficiency of genetic transfer, selection and regeneration of transformants (Birch, 1997).

*In vitro* regeneration of cucumber can be achieved by using different tissue culture approaches (Malepszy, 1988). Regeneration via organogenesis and somatic embryogenesis in this species has been described using a variety of explants, namely cotyledons (Trulson and Shahin, 1986; Chee, 1990; Gambley and Dodd, 1990; Punja et al., 1990a; Ali et al., 1991; Lou and Kako. 1994; Selvaraj et al., 2007; Grozeva and Velkov, 2014), hypocotyls (Ziv and Gadasi, 1986; Chee, 1990; Selvaraj et al., 2006; Grozeva and Velkov, 2014), leaves (Malepszy and Nadolska-Orczyk, 1983; Punja et al. 1990a; Lou and Kako. 1994; Burza and Malepszy, 1995a; Seo et al., 2000), and petiols (Punja et al. 1990a). Regeneration of cucumber has also been reported from protoplasts (Trulson and Shahin, 1986; Punja et al., 1990b; Burza and Malepszy. 1995b), and suspension cultures (Chee and Tricoli, 1988: Raharjo and Punja, 1994; Kreuger et al., 1996). Cucumber regeneration is largely dependent on genotype, explant type, and culture medium (Wehner and Locy, 1981; Trulson and Shahin, 1986; Punja et al. 1990a; Grozeva and Velkov, 2014; Li et al., 2016). Although *in vitro* plant regeneration of cucumber has been extensively reported, a high regeneration system for this species is still below the expected (Wang et al., 2015).

Inorganic macro- and micronutrients levels widely used in plant tissue culture media are based on those of Murashige and Skoog (1962) (MS) medium, developed for tobacco. However, for many of its micronutrients a clear optimal level was not evident. Furthermore, tissue culture of non-tobacco species may require a different concentration of its micronutrients for optimal response (Dahleen, 1995). Heavy metals such as copper (Cu), iron (Fe), manganese (Mn), molybdenum (Mo), nickel (Ni), and zinc (Zn) are essential plant micronutrients that are required in small but critical amounts for its normal growth and development (Alloway, 2013; Arif et al., 2016).

Cu is a redox-active transition element that is involved in many physiological processes in plants (Yruela, 2005). Its role as redox agent also makes it potentially toxic, as Cu ions can catalyse the production of damaging free radicals (Tomar et al., 2015). Cu acts as a structural component of numerous proteins and as a cofactor for many enzymes, including plastocyanin, cytochrome c oxidase, ascorbate oxidase, laccase, Cu/Zn superoxide dismutases (SODs), polyphenol oxidases, copper-containing amine oxidases, and phytohormone receptors. Cu plays a role in a wide range of processes such as photosynthetic electron transport, mitochondrial respiration, carbohydrate and nitrogen metabolism, cell wall metabolism, oxidative stress response, disease resistance, pollen viability, water permeability of xylem vessels, and hormone signaling (for reviews, see Yruela, 2009; Cohu and Pilon, 2010; Kabata-Pendias, 2010; Broadley et al., 2012; Tripathi et al., 2015; Migocka and Malas, 2018). Either deficient or in excess, it can cause disorders in plant growth and development (Yruela, 2009). Its deficiency leads to defects in photosynthesis, reduced respiration, chlorosis, and wilting of leaves (Shahbaz and Pilon, 2019), among others. Due to its relatively low mobility in plants, young organs are usually the first to develop Cu-deficiency symptoms (Kabata-Pendias, 2010). Cu levels in cells should be kept low, since this element is extremely toxic (due to its high redox properties) (Yruela, 2009). The most common symptoms of its toxicity are low photosynthesis efficiency, damage to DNA, disturbed protein complexes, damage to membrane permeability, chlorosis, and root malformation (Kabata-Pendias and Szteke, 2015).

It has been reported by different authors a positive effect of higher levels of copper in *in vitro* culture and morphogenesis of different plant species, namely in melon (Garcia-Sogo et al., 1991), watermelon (Ellul, 2002), carrot (Kowalska et al., 2012), indian ginseng (Sinha et al., 2010), tobacco (Purnhauser and Gyulai, 1993), tetraploid wheat (Ghaemi et al., 1994), hexaploid wheat (Purnhauser and Gyulai, 1993), triticale (Purnhauser and Gyulai, 1993), barley (Dahleen, 1995), indica rice (Sahrawat and Chand, 1999), sorghum (Nirwan and Kothari, 2003), bamboo (Singh et al., 2017), and date palm (Al-Mayahi, 2014). Nonetheless, negative effects of excessive exposure to this micronutrient have also been reported (e.g. Garcia-Sogo et al. (1991) in melon, Purnhauser and Gyulai (1993) in tobacco, and Ghaemi et al. (1994) in tetraploid wheat). To our knowledge, this is the first report on the effect of high concentration of copper ion on *in vitro* adventitious regeneration of cucumber.

The aim of the present work was to evaluate the influence of different concentrations of CuSO_4_ on *in vitro* adventitious organogenesis, using cotyledon as explants from one inbreed line and two commercial cultivars of cucumber.

## 2. Materials and Methods

### 2.1. Plant material and in vitro regeneration

Cucumber seeds from inbred line ‘Wisconsin 2843’ (kindly provided by Dr. C.E. Peterson, Michigan State University, East Lansing, USA), and cultivars ‘Marketer’ and ‘Negrito’ (Semillas Fitó S.A.) were used as starting material. Mature seeds were decoated and surface-sterilized by immersion in a diluted commercial bleach (5% w/v sodium hypochlorite) with 0.1 (v/v) 7X-O-matic (Flow Laboratories) for 30 min. The seeds were then rinsed three times with sterile distilled water for 5, 10 and 15 min, respectively. After sterilization, the seeds were germinated in the dark (150 mm x 25 mm test tubes) on solid germination medium (GM), consisting of Murashige and Skoog’s (1962) inorganic basal salts supplemented with 1% (w/v) sucrose, and 0.8% (w/v) agar (Industrial, Pronadisa), and its pH adjusted to 5.7 prior to autoclaving at 115 °C for 30 min. After 1 to 2 days, when the radicle emerged and curved into the medium, the test tubes were transferred to a tissue culture chamber maintained at 26 ± 2 °C with a 16 h photoperiod under cool-white fluorescent lamps (Grolux, Sylvania), and with a light intensity of 2000 lux. The same incubation conditions were used in the subsequent *in vitro* culture steps. Cotyledons from four-day-old seedlings were used as explant source by excising transversely 1-2 mm beyond its proximal and distal ends. The experimental assessments are based on naked-eye observations.

Cotiledonary explants were cultured for 3 weeks in 300 ml glass jars with the abaxial side in contact with the shoot induction medium (SIM), containing Murashige and Skoog’s (1962) inorganic basal salts supplemented with 3% (w/v) sucrose, 0.1 g L^−1^ myo-inositol, 1 mg L^−1^ thiamine-HCl, RT vitamins (Staba, 1969), 0.5 mg L^−1^ Indole-3-acetic acid (IAA), 2.5 mg L^−1^ 6-benzylaminopurine (BAP), and copper sulfate as CuSO_4_·5H_2_O (0.0, 0.2, 1.0 and 5.0 g L^−1^), in addition to its standard concentration of MS (0.025 mg L^−1^). The medium was solidified with 0.8% (w/v) agar (Industrial, Pronadisa) and its pH adjusted to 5.7 before autoclaving. At the end of this phase, the evaluation on callus regeneration frequency (%) (CRF) and callus extension index (CEI) was performed. The CRF (mean±SE) was calculated by the frequency of explants with callus formation in the cutting zone (proximal and distal edges); the CEI (mean±SE) was measured by the mean value corresponding to arbitrary values (from 0 to 3) on the extension of callus formation for each explant, whereas: 0 = absence of callus on the cutting zone; 1= traces of callus on the cutting zone; 2 = callus on less than half of the cutting zone; 3 = callus on half or more of the cutting zone; 4 = callus covering the full extension of the cutting zone.

Adventious buds and shoot primordia were then transferred to 300 ml glass jars containing shoot development and elongation medium (SDM) where auxins were not included and BAP was substituted by 0.2 mg L^−1^ KIN. After 2 weeks of culture, the response on shoot regeneration frequency (SRF) and shoot number index (SNI) was determined. The SRF (mean±SE) was calculated by the frequency of explants with shoot formation; the SNI (mean±SE) was measured by the mean value corresponding to arbitrary values (from 0 to 3) on the number of shoot formation for each explant, whereas: 0 = absence of any shoot; 1 = one shoot; 2 = 2 shoots; 3 =3 or more shoots. Individualized shoots were further rooted on hormone-free MS medium and within 3 to 4 weeks, the plantlets were ready for acclimatization (data not shown).

### 2.2. Data analysis

The experimental design was a two-way factorial arrangement 3 × 4 on a completely randomized design, with at least 11 replicate jars per treatment with 6 explants per jar. Treatment means comparison were assessed by non-linear regression analyses. Count data variables, i.e. callus extension index (CEI) and shoot number index (SNI) were submitted to generalized poisson regression and negative binomial regression analyses, respectively. Logistic regression was performed for data collected as binary values. Relationships between SNI (response variable) and the predictors (CEI, CuSO_4_ concentration) were analyzed using generalized poisson regression. All statistics were performed using R 3.4.4 software (R Core Team, 2018). The R packages ‘stats’ (R Core Team, 2018), ‘VGAM’ (Yee, 2019), and ‘MASS’ (Venables and Ripley, 2002) were used to perform logistic regression, negative binomial regression, and generalized poisson regression, respectively. Dispersion test was performed using the ‘AER’ package (Kleiber and Zeileis, 2008), in order to test the suitability of applying Poisson GLMs to our count data. Binary data were tested for overdispersion by fitting the model twice, i.e., firstly using a binomial family, and then a quasibinomial family, and by using the ‘pchisq’ function (Kabacoff, 2011) from the ‘stats’ package, indicating a lack of overdispersion. Multicollinearity between predictors was assessed by calculating the generalized variation inflation factor (GVIF; Fox and Monette, 1992) for each predictor, using the ‘vif’ function in the ‘car’ package (Fox and Weisberg, 2011), indicating that multicollinearity was not a problem in the present analyses (GVIFs<1.24). The ‘rockchalk’ package (Johnson, 2019) was used to combine levels not significantly different from each other in the analysis of the association between variables. Model performance was evaluated using Akaike Information Criterion (AIC; Akaike, 1973) and Bayesian Information Criterion (BIC; Schwarz, 1978), and the best fitting model was selected based on the lowest AIC and BIC values. The Wald test was used in regression to test for significance. A probability value of P < 0.05 was used as the criterion for statistical significance for all analyses.

## 3. Results

### 3.1. Frequency and extension of callus

The results are shown in Table 1. Statistics numerical data are expressed as mean ± standard error of the mean (SEM). The formation of callus occurred in the first 2 weeks of culture, starting from the cut edges of the primary explants. All treatments recorded 100% callus regeneration frequency. Data collected as arbitrary values were submitted to regression analysis. Callus extension index (CEI) showed significant differences in two of the three cultivars of cucumber for ‘CuSO_4_ concentration’ (P < 0.05). The response in CEI, ranked from highest to lowest, varied as follows: for ‘Marketer’, from 2.60±0.06 (0.2 mg L-1 CuSO_4_) to 2.91±0.05 (0 mg L-1 CuSO_4_); for ‘Negrito’, with no significant differences, from 2.52±0.07 (1.0 mg L-1 CuSO_4_) to 2.70±0.06 (0.2 mg L-1 CuSO_4_); and for ‘Wisconsin 2843’, from 2.03±0.02 (5.0 mg L-1 CuSO_4_) to 2.52±0.06 (1.0 mg L-1 CuSO_4_). These results are between the arbitrary values of 2: callus on less than half of the cutting zone, and 3: callus on half or more of the cutting zone. In virtually all cases, the order of magnitude of the CEI response of cultivars to the CuSO_4_ concentration was different. In the control group, callus of the ‘Wisconsin 2843’ line contrasts with the lighter and more friable callus of the ‘Marketer’ and ‘Negrito’ cultivars. With the addition of high concentrations of CuSO_4_, the callus generally became more compact and greenish (data not shown).

**Table 1.**
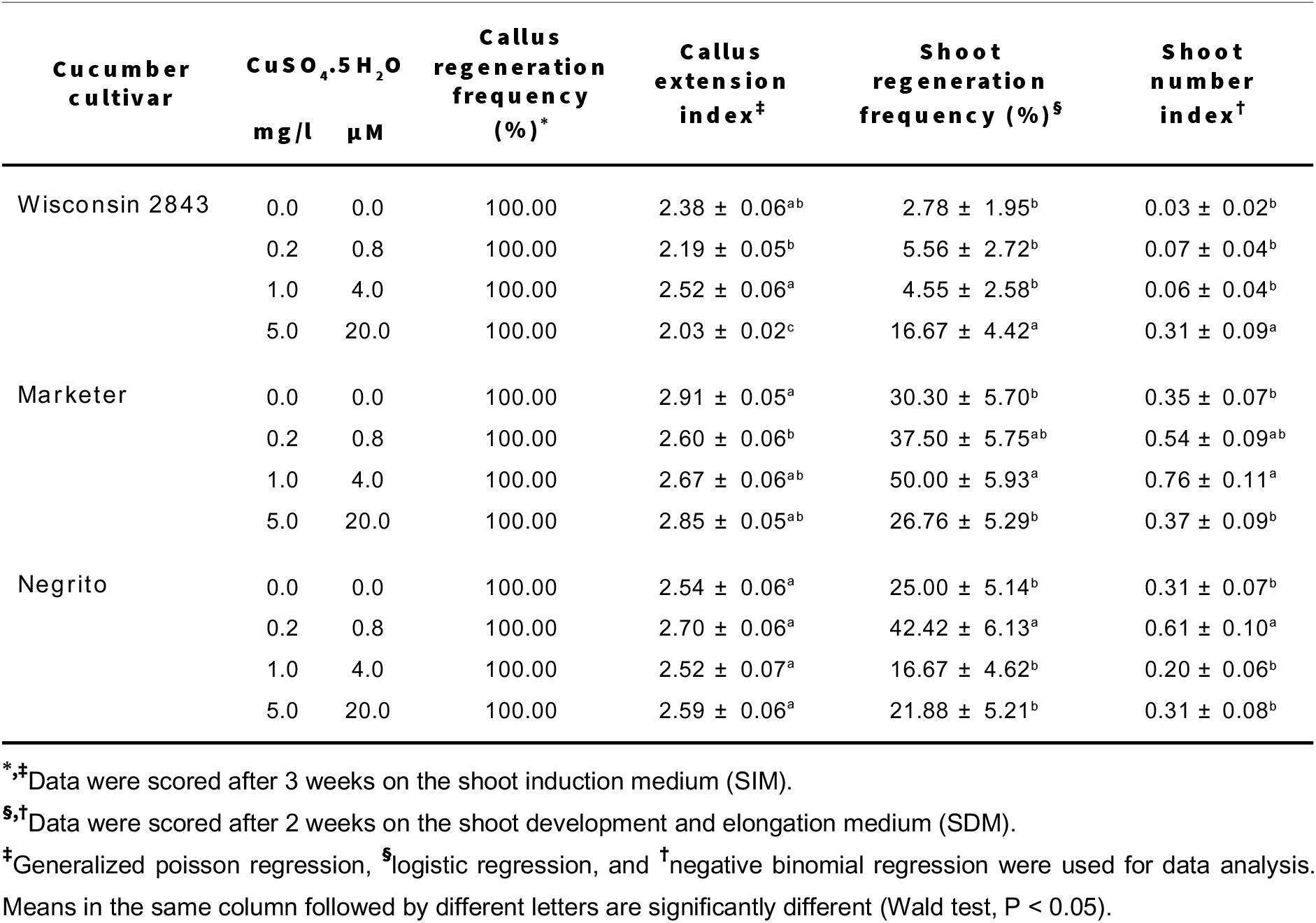
Effect of the addition of different concentrations of copper sulfate to MS-derived shoot induction medium (SIM) containing 0.5 mg L-1 IAA and 2.5 mg L-1 BAP on *in vitro* callus and shoot regeneration from cotyledon explants of three cultivars of *Cucumis sativus* L. Values are expressed as mean ± standard error of the mean.

### 3.2. Frequency and number of shoots

Results are presented as mean ± SEM (Table 1). Significance was set at P < 0.05. At 1.5-3 weeks of culture, groups of adventitious buds emerge especially at the proximal end of the explant, which at 2-4 weeks under cultivation could give rise to shoot primordia. Binary and count-data variables, i.e. shoot regeneration frequency (%) (SRF) and shoot number index (SNI), respectively, were subjected to regression analysis. Both variables revealed significant differences in all the three cultivars of cucumber for ‘CuSO_4_ concentration’. For ‘Wisconsin 2843’, both SRF and SNI tended to increase with increasing concentration of CuSO_4_; the highest SRF (16.67±4.42) and SNI (0.31±0.09) was obtained for the medium supplemented with 5.0 mg L-1 CuSO_4_, resulting in 6- and 10.3-fold higher, respectively, than the control. In the case of ‘Marketer’ both variables increased with increasing concentration of CuSO_4_ up to 1.0 mg L-1, but thereafter decreased significantly as the concentration increases; the maximum SRF (50.00±5.93) and SNI (0.76±0.11) resulted in 1.7- and 2.2-fold greater, respectively, when compared to the control. In relation to ‘Negrito’ both variables increased with 0.2 mg L-1 of CuSO_4_, but thereafter decreased significantly as the concentration increases; the highest SRF (42.42±6.13) and SNI (0.61±0.10), resulted in 1.7- and 2-fold higher, respectively, than the control. All cultivars revealed significant differences between the medium with the highest response and the control. From the above results, the performance of the cultivars, ranked from highest to lowest, was as follows: ‘Marketer’>‘Negrito’>‘Wisconsin 2843’.

### 3.3. Relationship between variables

The results are given in Table 2. The relationships between the variables were assessed through multiple regression analysis, with shoot number index (SNI) as the response variable, and callus extension index (CEI) and ‘CuSO_4_ concentration’ as the predictors. Statistical significance was defined as P < 0.05. No relationship was observed between SNI and CEI in all the three cucumber cultivars tested. On the contrary, a strong relationship was found between SNI and ‘CuSO_4_ concentration’ (Wisconsin 2843: P=0.0027; Marketer: P=0.0016; Negrito: P=0.0002).

**Table 2.**
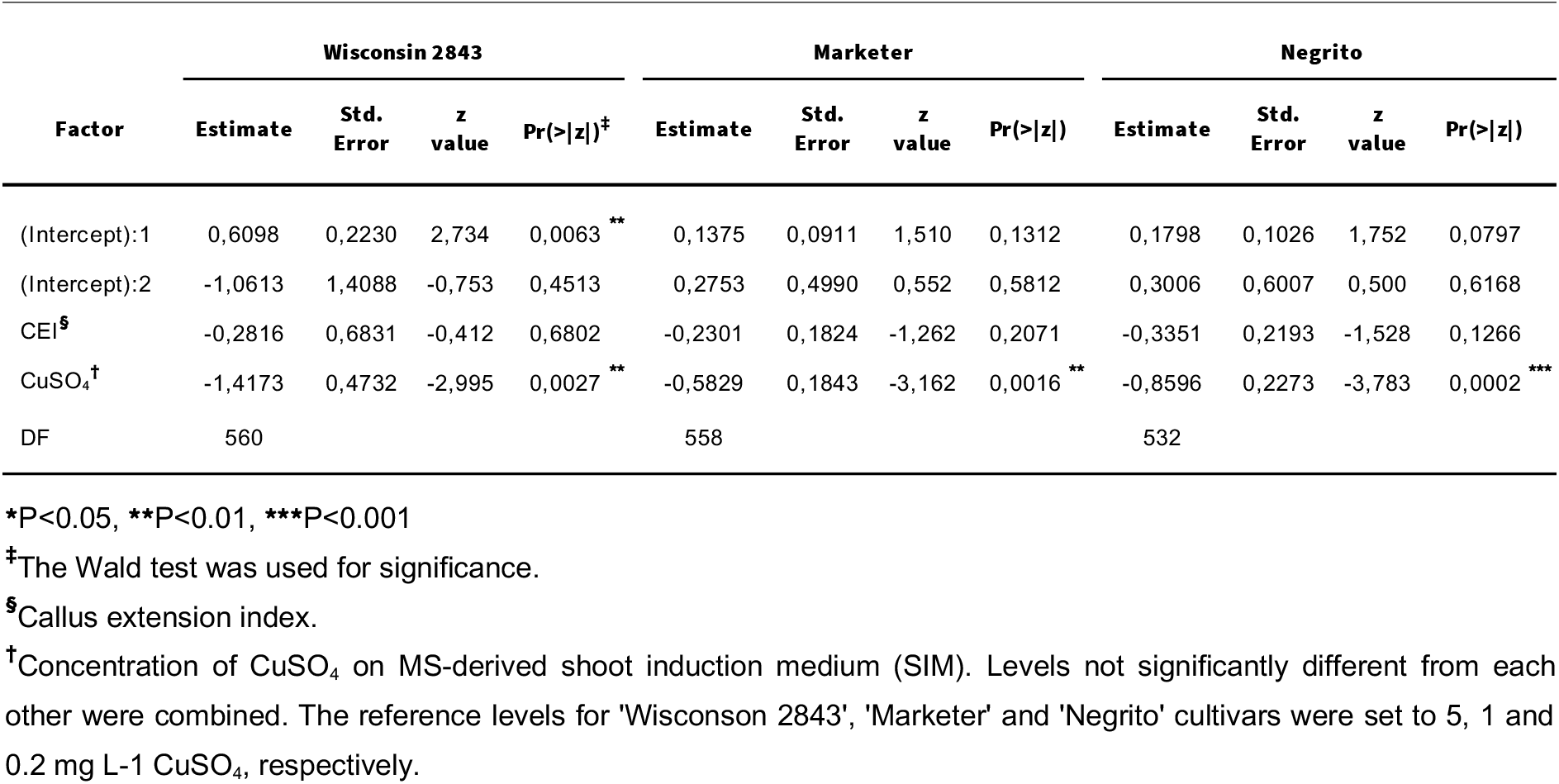
Factors associated with shoot number index (SNI) in generalized poisson regression among three cultivars.

## 4. Discussion

The purpose of this work was to evaluate the effect of different concentrations of copper sulfate on the organogenic response of several cucumber cultivars. Cooper is an essential plant micronutrient that is required at low concentrations. For instance, in the Murashige and Skoog’s (1962) mineral solution, copper is added as copper sulfate at a concentration of 0.025 mg L-1. Different studies on *in vitro* plant regeneration showed that MS basal medium could be optimized for a specific genotype by adjusting the concentration of some of its micronutrients (e.g. Dahleen, 1995; Sahrawat and Chand, 1999; Kothari-Chajer et al., 2008).

Most essential heavy metals are required in plants and animals for biochemical and physiological functions. They integrate various key enzymes and have major roles in different oxidation-reduction reactions (Nunes and Malmlöf, 2018). Heavy metals such as copper (Cu), iron (Fe), manganese (Mn), molybdenum (Mo), nickel (Ni), and zinc (Zn) are essential plant micronutrients that are required in very small amounts for its normal growth and development. Excessive levels of these elements within the plant tissue have different toxic effects, such as oxidative stress with further inhibition of cytoplasmic enzymes and damage to cellular structures, and a change on the uptake and accumulation of essential to non-essential elements, which affects enzymes and proteins function, and metabolisms (Arif et al., 2016).

In the present study, we observed that all explants formed callus and, the response on callus extension was dependent on the genotype and, in two of the three cultivars on the level of copper sulfate in the medium. Callus regeneration was usually observed in both proximal and distal zones of the cotyledon whereas shoot-buds were formed almost exclusively from the proximal part. This is in general agreement with other studies in cucurbits and other plants in which the proximal part of the cotyledon, adjacent to the cut edge of the explant, is the most active site for the regeneration of organized structures: cucumber (Gambley and Dodd, 1990; Msikita et al., 1990); melon (Gaba et al., 1999); horned melon (Lin et al., 2011); watermelon (Compton and Gray, 1993); squash (Ananthakrishnan et al., 2003); bottle gourd (Han et al., 2004); soybean (Hinchee et al., 1988); geranium (Chang et al., 1996). Cotyledon polarity on adventitious shoot regeneration is most likely the key factor for the absence of association between the number of shoots and the extension of callus, obtained for all cultivars.

The results on shoots regeneration showed dependence on the genotype and on the level of copper sulfate in the medium, indicating a possible interaction between the two factors. For both frequency (SRF) and number of shoots (SNI) the optimal concentration of CuSO_4_ was 8-200 fold higher than the control (0.025 mg L^−1^ - the standard MS concentration) and was cultivar-specific. Differences in the ability for callus formation and regeneration have been extensively reported among cucumber genotypes (e.g. Wehner and Locy, 1981; Grozeva and Velkov, 2014; see mini-review by Wang et al., 2015), and within other species. Likewise, when high concentrations of copper were applied, differences have been reported within rice (Sahrawat and Chand, 1999), and barley (Dahleen, 1995), with the optimal concentration of the ion being cultivar-specific.

The positive effects of high concentrations of copper sulfate on *in vitro* adventitious shoot organogenesis of cucumber, are in line with those obtained with other cucurbits. Garcia-Sogo et al. (1991) found that levels of CuSO_4_ ranging from 0.1 to 5 mg L^−1^ on MS-based culture media resulted in a significant increase in callus-derived shoot-buds from cotyledon explants of melon. Ellul (2002) observed a positive response on the regeneration of watermelon from cotyledon explants when concentrations of CuSO_4_ ranging from 0.1 to 1.0 mg L^−1^ on MS-based culture media were used. The beneficial effects of high concentrations of copper on *in vitro* morphogenesis and regeneration are not exclusive to cucurbits. Ghaemi et al. (1994) found that levels of CuSO_4_ ranging from 2 to 10 mg L^−1^ increased the production of embryoids from anthers in three of the four tetraploid wheat genotypes tested. Other studies reported that concentrations of CuSO_4_ ranging from 0.125 to 25 mg L-1 significantly enhanced shoot and root regeneration in hexaploid wheat and triticale callus cultures initiated from immature embryos, shoot regeneration from leaf-disc cultures of tobacco (Purnhauser, 1991; Purnhauser and Gyulai, 1993), and plant regeneration from barley callus cultures derived from immature embryos (Dahleen, 1995). Callus growth and shoot regeneration in date palm callus cultures initiated from shoot apical meristem explants were significantly enhanced with 0.5 mg L^−1^ CuSO_4_, and root regeneration by using CuSO_4_ at 0.125 mg L-1 (Al-Mayahi, 2014). Optimal shoot regeneration from nodal explants of bamboo was achieved with the addition of CuSO_4_ five-fold higher than in MS medium (Singh et al., 2017). The enhancement of adventitious shoot formation with the application of high concentrations of copper sulfate was accompanied by an increase in the qualitative response, i.e., greener and more vigorous shoots (data not shown). Likewise, the presence of high levels of CuSO_4_ on adventitious regeneration of melon (Souza et al., 2006), finger (Kothari et al., 2004; Kothari-Chajer et al., 2008) and kodo millets (Kothari-Chajer et al., 2008), improved the quality of response.

It is known that some heavy metals play important roles in the regeneration of plant tissue culture (Purnhauser and Gyulai, 1993). Besides Cu, other metals such as Ag, Co, Mn, Ni, and Zn have been also reported to stimulate morphogenesis (Roustan et al., 1989; Purnhauser and Gyulai, 1993; Kothari et al., 2004; Kothari-Chajer et al., 2008).

The basis of the positive effect of copper in plant morphogenesis is still not clear. According to Purnhauser and Gyulai (1993) some Cu enzymes might play a major role in plant regeneration, since Cu^2+^ is a component or activator of many important enzymes that participate in electron transport, protein and carbohydrate biosynthesis, polyphenol metabolism, and so forth. Our results on the effect of copper on callus extension and on the number and development of organized structures (adventitious buds and shoots), may suggest that this metal ion could modulate genes related to regeneration and/or alter the level or activity of endogenous growth regulators. Furthermore, adverse effects generated by *in vitro* culture, such as those caused by excessive ethylene, reactive oxygen species (ROS), and (poly) phenols, could be mitigated by increased copper levels. Other studies may indirectly suggest the importance of copper in plant morphogenesis. The respiration rate in cells under callus proliferation and cell division is normally higher (Al-Mayahi, 2014), and enhanced levels of copper may be required, since it plays a key role in the respiration process (Yruela, 2009). Rapid cell division and differentiation require adequate amounts of precursors for cell wall biosynthesis (Prażak and Molas, 2015), where copper metalloenzymes play an important part (Delhaize et al., 2015). The molecular and physiological mechanisms associated with this essential, but potentially toxic, heavy metal, should be further investigated to understand its effect on *in vitro* plant regeneration.

## 5. Conclusions

To the best of our knowledge, this is the first report on the effect of high concentrations of copper ion on *in vitro* adventitious regeneration of cucumber. From this study, we conclude that optimized copper sulfate levels in MS basal medium, effectively enhanced shoot organogenesis in one inbred line and two commercial cultivars of cucumber. These findings can be further used for large-scale micropropagation of this species, and in genetic transformation studies, could help improve the regeneration efficiency of transformed cells.

## Conflict of interests

The author declare that there is no conflict of interests regarding the publication of this paper.

## Acknowledgements

The Author is grateful to the Spanish Agency for International Development Cooperation (Agencia Española de Cooperación Internacional para el Desarrollo, AECID) (Spanish Government) for the PhD fellowship.

## Notes

### Competing Interest Statement

The authors have declared no competing interest.

